# Comprehensive analysis of prognostic immune-related genes in the tumor microenvironment of lung adenocarcinoma

**DOI:** 10.1101/2020.03.09.983080

**Authors:** Chao Ma, Huan Luo, Jing Cao

**Affiliations:** Charité – Universitätsmedizin Berlin, corporate member of Freie Universität Berlin, Humboldt-Universität zu Berlin, and the Berlin Institute of Health; Charité - Universitätsmedizin Berlin, BCRT - Berlin Institute of Health Center for Regenerative Therapies, Berlin, Germany; Klinik für Augenheilkunde, Charité – Universitätsmedizin Berlin, Corporate Member of Freie Universität Berlin, Humboldt-Universität zu Berlin, and Berlin Institute of Health, Berlin, Germany; Department of Human Anatomy, School of Basic Medicine, Zhengzhou University, Zhengzhou, China

**Keywords:** TCGA, GEO, tumor microenvironment, immune scores, stromal scores, overall survival, lung adenocarcinoma

## Abstract

**Background:** Lung adenocarcinoma (LUAD) is the most common primary lung cancer, and increasing evidence indicates the clinical importance of the microenvironment in LUAD. Tumor-infiltrating immune cells play an important role in promoting or inhibiting tumor growth. This study aimed to identify immune-related prognostic genes that were associated with the LUAD microenvironment.

**Methods:** We used the “estimate” R package to calculate the immune/stromal scores of each sample of GSE72094 based on the ESTIMATE algorithm. Then we looked up relationships between patients’ characteristics and immune/stromal scores. After that, we divided the samples into two groups: high and low scores, identified the common differentially expressed genes (DEGs), and performed Gene Ontology (GO) and Kyoto Encyclopedia of Genes and Genomes (KEGG) on the common DEGs. After conducting the overall survival analysis of the common DEGs, prognostic genes were harvested. Then we constructed the protein-protein interaction network and performed the enrichment of GO and KEGG for the prognostic genes. Crucial prognostic genes were obtained after validating in two independent data sources (GSE68465 and TIMER). Finally, we investigated the immune correlates of the crucial prognostic genes based on the TIMER.

**Results:** Immune scores did not vary with gender, age, smoking history, tumor stage, and EGFR status, but vary with the status of KRAS, STK11, and TP53. For the stromal scores, only the status of STK11 and TP53 mattered. Reduced immune score predicted poor prognosis of LUAD. 357 common DEGs were found, of which 108 were identified as prognostic genes after overall survival analysis. GO and KEGG analysis found that common DEGs and prognostic genes were both mainly involved in immune-related items. After validation in two independent data sources, 12 genes were validated to be crucial prognostic genes linked to prognosis. After investigated the TIMER, all 12 genes were correlated with the main immune cell types.

**Conclusion:** 12 immune-related prognostic genes were discovered relating to the microenvironment in LUAD. These findings suggest that the composition of the tumor microenvironment affects the clinical outcomes of LUAD, and it may provide a basis for the development of novel prognostic biomarkers and immunotherapy for LUAD.

## Introduction

Lung cancer occurred in approximately 2.1 million patients in 2018 and caused an estimated 1.7 million deaths worldwide^1^. In the United States, there will be approximately 230,000 new cases of lung cancer, and over 140,000 deaths annually^2^. Lung cancer is a group of heterogeneous tumors consisting of more than 50 histomorphological subtypes^3^. In the past few decades, the two diagnostic terms, non-small cell lung cancer (NSCLC) and small cell lung cancer, have been used most frequently^4^. Lung adenocarcinoma (LUAD) is the most diagnosed histological subtype of NSCLC, followed by squamous cell carcinoma^1–3^.

Advances in understanding the molecular pathogenesis of LUAD could provide important information regarding prognosis^5^. The environment in which the tumor exists is called the tumor microenvironment and is composed of fluids, extracellular matrix molecules, immune cells, stromal cells, and inflammatory cells^6^. The tumor microenvironment is in a highly dynamic, its importance in jointly promoting the immune escape, growth, and metastasis of tumors has been well recognized, of which reflecting the evolutionary nature of cancer^6,7^. For example, levels of immune cell infiltration have been reported to be related to prognosis, and the activity of both immune and stromal cells has been shown to predict overall cancer survival^8^. Each component of the microenvironment plays an important role in LUAD, and therefore has a significant impact on clinical outcomes^5,9–15^.

In 2013, Yoshihara et al. designed an algorithm called ESTIMATE (Estimation of STromal and Immune cells in MAlignant Tumor tissues using Expression data). The algorithm analyzes specific gene expression characteristics of immune and stromal cells and calculates immune and stromal scores to predict non-tumor cell infiltration^16^. With the help of the ESTIMATE algorithm, researchers conducted prognostic evaluations and exploration of genetic changes in many malignancies^17–19^. However, the immune and stromal scores of LUAD remain to be elucidated. It is unclear whether the ESTIMATE algorithm can be used to investigate the prognosis of LUAD. Previous studies have shown that immune cell infiltration is associated with prognosis, which means that assessing the heterogeneity of the tumor microenvironment pattern and reshaping the immune microenvironment may hold promise for future cancer treatments^8,20,21^. Assessing the immune cell infiltration of genes may provide more insights for immunotherapy.

For better understanding the molecular pathogenesis of LUAD, in the present work, we applied the ESTIMATE and Tumor IMmune Estimation Resource (TIMER) algorithm, and three datasets (one for modeling and two for verification) to identify the crucial immune-related prognostic genes of LUAD.

## Materials and Methods

### Data collection

We searched the Gene Expression Omnibus (GEO, http://www.ncbi.nlm.nih.gov/geo/) database (by using “lung adenocarcinoma” as the keywords. Then we filtered the results by checking “Series” as the “Entry type”, “Expression profiling by array” as the “Study type”, and “Homo sapiens” as the “Organism”. In the filtered results, we delve into the details of each dataset and select only the dataset with more than 400 samples that contained survival data. Finally, datasets GSE72094 and GSE68465 (**Table 1**) were selected for this study, and the expression and clinical data of them were downloaded. In this study, GSE72094 was used to build the model, while, GSE68465 was applied to validate it.

**Table 1.** Basic information of GEO datasets included in this study.

| Series | Year | Country | Platform | Cases |
| --- | --- | --- | --- | --- |
| GSE7209<br>4 | 201<br>5 | USA | GPL15048, Rosetta/Merck Human RSTA Custom Affymetrix 2.0 microarray [HuRSTA_2a520709.CDF] | 442 |
| GSE6846<br>5 | 201<br>5 | USA | GPL96, [HG-U133A] Affymetrix Human Genome U133A Array | 462 |

GEO: Gene Expression Omnibus

### Calculation of immune and stromal scores

ESTIMATE is a tool for predicting tumor purity, and the presence of infiltrating stromal/immune cells in tumor tissues using gene expression data^16^. To begin with, “estimate” R package was applied to calculate immune and stromal scores based on the gene expression data of GSE72094.

### Relationships between patients’ characteristics and immune/stromal scores

All cases were grouped according to their clinical or pathological features. One-way ANOVA test was utilized to assess the relationship between the characteristics and immune/stromal scores. P-value < 0.05 was considered statistically significant. Overall survival was applied as the primary prognostic endpoint, which was estimated by the Kaplan-Meier survival estimator. Patients were then stratified according to the median of immune and stromal, with high and low scores. “survival” R package was applied to perform overall survival analysis between high and low score groups. The overall survival of the two groups were compared by log-rank tests. P-value < 0.05 was considered statistically significant.

### Differentially expressed gene analysis

Differentially expressed genes (DEGs) were identified between high and low immune/stromal score groups using “limma” R package^22^. Genes with |log_2_(fold-change) | > 0.75 and false discovery rate (FDR) < 0.05 were selected as DEGs. “pheatmap” R package was applied to produce heatmaps and clustering of DEGs.

### Enrichment analysis of common DEGs

Among the above comparison, genes who upregulated/downregulated in both high immune and stromal score groups were identified as common upregulated/downregulated DEGs. Common DEGs were a combination of them. Functional enrichment analysis of common DEGs, including gene ontology (GO) and the Kyoto Encyclopedia of Genes and Genomes (KEGG), were performed using “clusterProfiler” R package. FDR < 0.05 was considered statistically significant.

### Identification of prognostic genes

Kaplan-Meier plots were generated using the “survival” R software package, which illustrates the relationship between overall survival and the expression level of each common DEG gene. The relationship was tested by a log-rank test. The gene with p-value < 0.05 was identified as the prognostic gene. Subsequently, prognostic genes were applied for functional enrichment analysis (FDR < 0.05).

### Establishment of the protein-protein interaction (PPI) network of prognostic genes

To deeply investigate prognostic genes, we built the PPI network of prognostic genes with the help of the STRING online tool (http://string-db.org) and Cytoscape software (http://www.cytoscape.org/). We import prognostic genes into the STRING database for calculation. Besides, filtered the results only by checking the options of “hiding disconnected nodes in the network”, leaving other settings default. Then the calculation result from STRING was imported to Cytoscape for visualization. The Cytoscape “cytoHubba” plug-in and Network Analyzer were applied to analyze the degree distribution. The Cytoscape “MCODE” plug-in was applied to find clusters (highly interconnected regions) in this PPI network.

### Validation in two independent data source

We validated the previously identified prognostic genes using dataset GSE68465 (462 LUAD patients included) by overall survival. Genes with p-value < 0.05 (Log-rank test) were considered significant. Tumor IMmune Estimation Resource (TIMER, https://cistrome.shinyapps.io/timer/) incorporated 10,009 samples across 23 cancer types (506 LUAD patients included) from The Cancer Genome Atlas (TCGA, https://portal.gdc.cancer.gov/). The survival module in it can be used for exploring the association between clinical outcome and gene expression^23,24^. Hence, we use TIMER to validate the prognostic genes in LUAD by overall survival, additionally. Genes with p-value < 0.05 (Log-rank test) were considered significant. The genes in the intersection of the results from GSE68465 and TIMER were identified as crucial prognostic genes.

### Validation of the immune correlates of the crucial prognostic genes

The TIMER also included data of abundances of 6 main immune infiltrates (B cells, CD4+ T cells, CD8+ T cells, Neutrophils, Macrophages, and Dendritic cells), which are estimated by TIMER statistical method and validated using pathological estimations^23,24^. Compared to other calculation methods, the TIMER can eliminate bias effects by removing highly expressed genes and eliminating collinearity between immune cells to ensure accuracy of the calculation^25^. The “Gene module” in TIMER can explore the correlation between gene expression and abundance of immune infiltrates in LUADs. We employed TIMER for validating the immune cell infiltration of crucial prognostic genes. The partial correlation coefficient and corresponding p-value are visualized via “canvasXpress” R package.

## Results

### Characteristics of microarray data and patients

Dataset GSE72094 containing 442 cases with microarray expression and clinical data from GEO database was analyzed. For details of the microarray data, see **Table 1**. Among the clinicopathological characteristics of patients, 240(54.3%) were females, 202(45.7%) were males. 60(13.57%) were younger than 60 years old and 361(81.67%) were equal or older to 60, while the rest were unknown. In the smoking history distribution, 335(75.79%), 33(7.47%), and 74(16.74%) were ever smoker, never smoker, and unknown, respectively. In tumor stage distribution, 265(59.95%) patients were in stage 1, 69(15.61%) were in stage 2, 63(14.25%) were in stage 3, 17(3.85%) were in stage 4, and 28(6.33%) were unknown. Genetic mutation information was also included. Mutations occurred in KRAS, EGFR, STK11, and TP53 accounted for 154 (34.84%), 47 (10.63%), 68 (15.38%), and 111 (25.11%), respectively. In the present study, 298(67.42%) patients were alive, 122(27.6%) were dead, 22(4.98%) were information unavailable (**Table 2**).

**Table 2.** The characteristics of patients based on the immune and stromal scores.

| Characteristic | Immune score<br>(range) | Stromal score<br>(range) | Cases<br>(percentage) |
| --- | --- | --- | --- |
| Gender |  |  |  |
| female | (-420.37~3609) | (-630.34~2227.47) | 240(54.3%) |
| male | (-266.91~3419.93) | (-1158.81~2412.18) | 202(45.7%) |
| Age |  |  |  |
| < 60 years old | (466.59~3609) | (-135.47~2227.47) | 60(13.57%) |
| ≥ 60 years old | (-420.37~3571.59) | (-1158.81~2412.18) | 361(81.67%) |
| unknown | (523.21~3341.4) | (-630.34~2048.95) | 21(4.75%) |
| Smoking history |  |  |  |
| ever | (-420.37~3609) | (-1158.81~2412.18) | 335(75.79%) |
| never | (718.12~3419.93) | (-135.47~1875.31) | 33(7.47%) |
| unknown | (520.17~3389.01) | (-630.34~2021.33) | 74(16.74%) |
| Tumor stage |  |  |  |
| 1 | (-266.91~3583.31) | (-1158.81~2412.18) | 265(59.95%) |
| 2 | (397.31~3419.93) | (-217.93~1988.13) | 69(15.61%) |
| 3 | (466.59~3609) | (-502.86~2227.47) | 63(14.25%) |
| 4 | (-420.37~3141.39) | (-740.24~2159.18) | 17(3.85%) |
| unknown | (523.21~3341.4) | (-630.34~2048.95) | 28(6.33%) |
| KRAS status |  |  |  |
| mutation | (-266.91~3609) | (-1158.81~2227.47) | 154(34.84%) |
| wild type | (-420.37~3583.31) | (-740.24~2412.18) | 288(65.16%) |
| EGFR status |  |  |  |
| mutation | (523.21~3419.93) | (-630.34~1940.25) | 47(10.63%) |
| wild type | (-420.37~3609) | (-1158.81~2412.18) | 395(89.37%) |
| STK11 status |  |  |  |
| mutation | (-266.91~3400.34) | (-1158.81~2226.55) | 68(15.38%) |
| wild type | (-420.37~3609) | (-815.95~2412.18) | 374(84.62%) |
| TP53 status |  |  |  |
| mutation | (-420.37~3609) | (-815.95~2227.47) | 111(25.11%) |
| wild type | (-266.91~3583.31) | (-1158.81~2412.18) | 331(74.89%) |
| Survival status |  |  |  |
| alive | (264.05~3609) | (-815.95~2412.18) | 298(67.42%) |
| dead | (-420.37~3333.28) | (-1158.81~2226.55) | 122(27.6%) |
| unknown | (523.21~3341.4) | (-630.34~2048.95) | 22(4.98%) |

### Relationships between patients’ characteristics and immune/stromal scores

We showed in detail in **Table 2** the distribution of immune / stromal scores at different clinicopathological characteristics. As shown in **Figure 1**, the distributions of immune scores did not vary with gender, age, smoking history, tumor stage, and EGFR status, but vary with the status of KRAS, STK11, and TP53. For the distributions of the stromal scores, only the status of STK11 and TP53 mattered.

**Figure 1.**
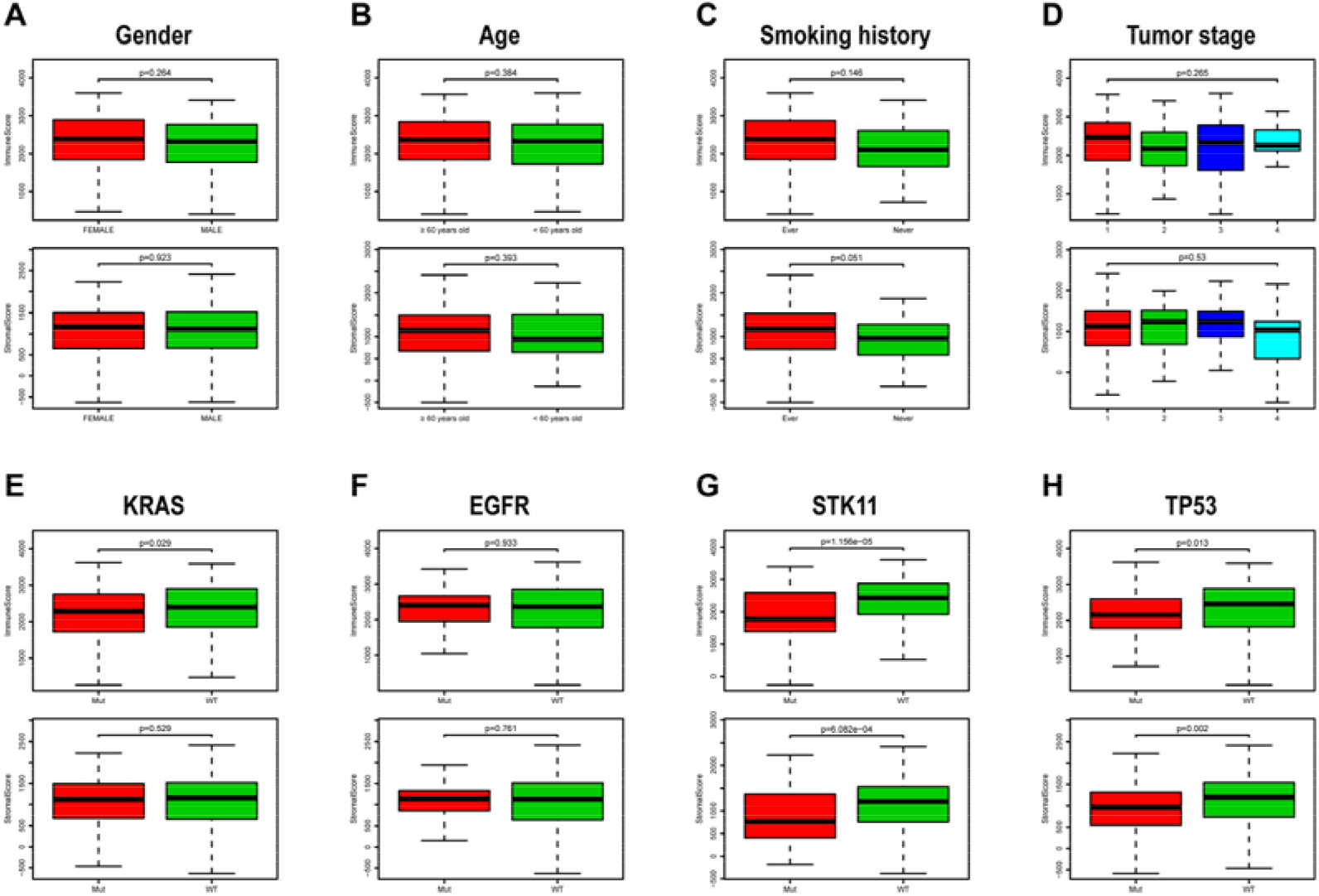
Relationships between patients’ characteristics and immune/stromal scores. Distribution of immune scores (**upper**) and stromal scores (**bottom**) plotted against gender (**A**), age (**B**), smoking history (**C**), tumor stage (**D**), KRAS status (**E**), EGFR status (**F**), STK11 status (**G**), and TP53 status (**H**). P-value < 0.05 was considered statistically significant. Mut: mutation; WT: wild type.

### Reduced immune score predicted a poor prognosis

To locate the potential correlation of overall survival with immune/stromal scores, we divided the 442 patients into high and low score groups according to their scores. Kaplan-Meier survival curves showed that the overall survival trend of patients in the high immune score group is more favorable than the cases in the low score group (p-value = 0.028 in log-rank test) (**Figure 2A**). As shown in **Figure 2B**, patients with high stromal scores achieved relatively good overall outcomes (p-value = 0.546 in log-rank test).

**Figure 2.**
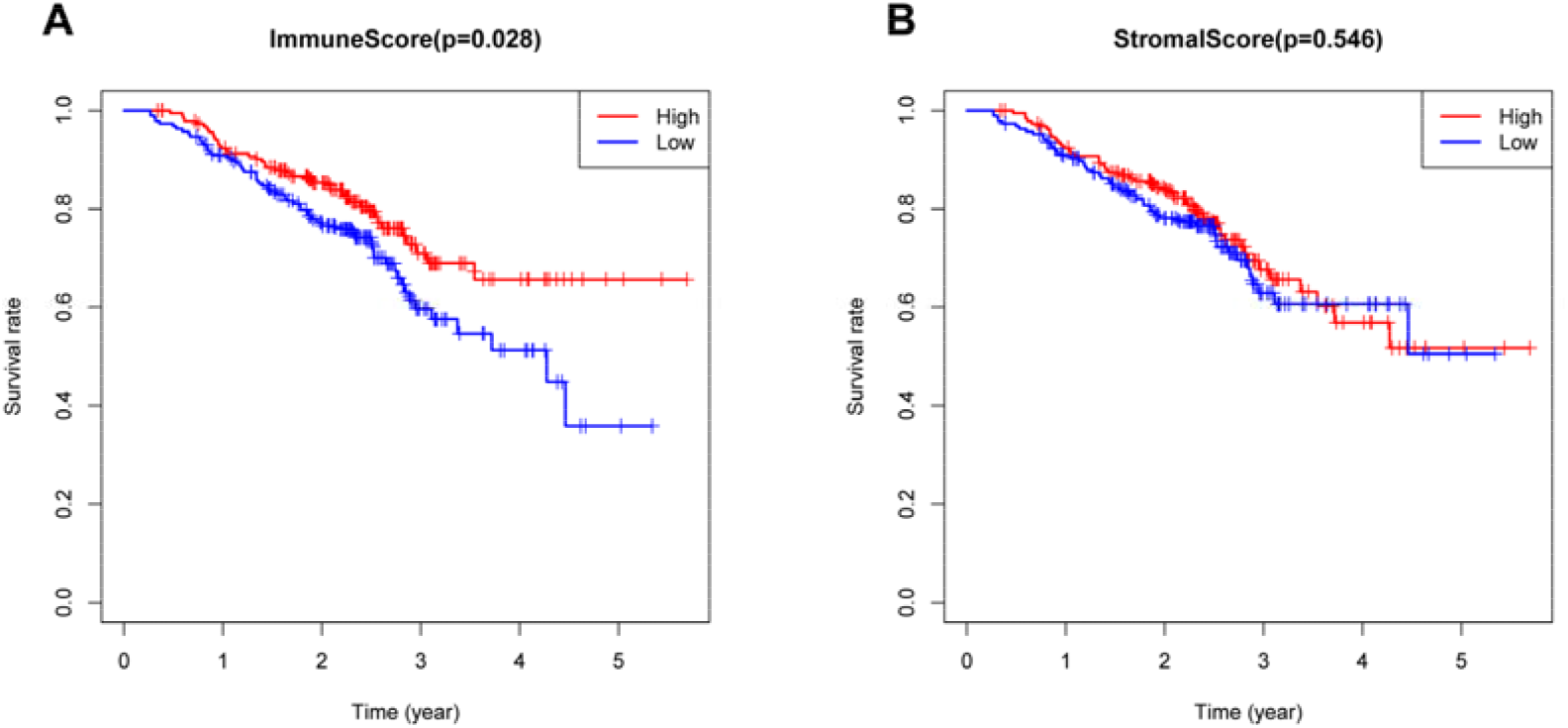
Kaplan-Meier overall survival curves based on immune/stromal scores. Patients were classified into high (**red**) and low (**blue**) immune (**A**) / stromal (**B**) scores groups.

### Comparison of gene expression profiles based on immune and stromal scores

In order to correlate gene expression profiles based on immune and stromal scores, the 442 cases were divided into groups of high and low scores according to their scores based on the median. **Figure 3A** shows a heatmap of 698 DEGs between immune score groups. **Figure 3B** displayed a heatmap consisting of 672 DEGs between stromal score groups. Via integrated bioinformatics analysis, we identified 348 common upregulated DEGs (**Figure 3C**) and 9 common downregulated (**Figure 3D**) DEG. Our subsequent analysis focused on these 357 common DEGs.

**Figure 3.**
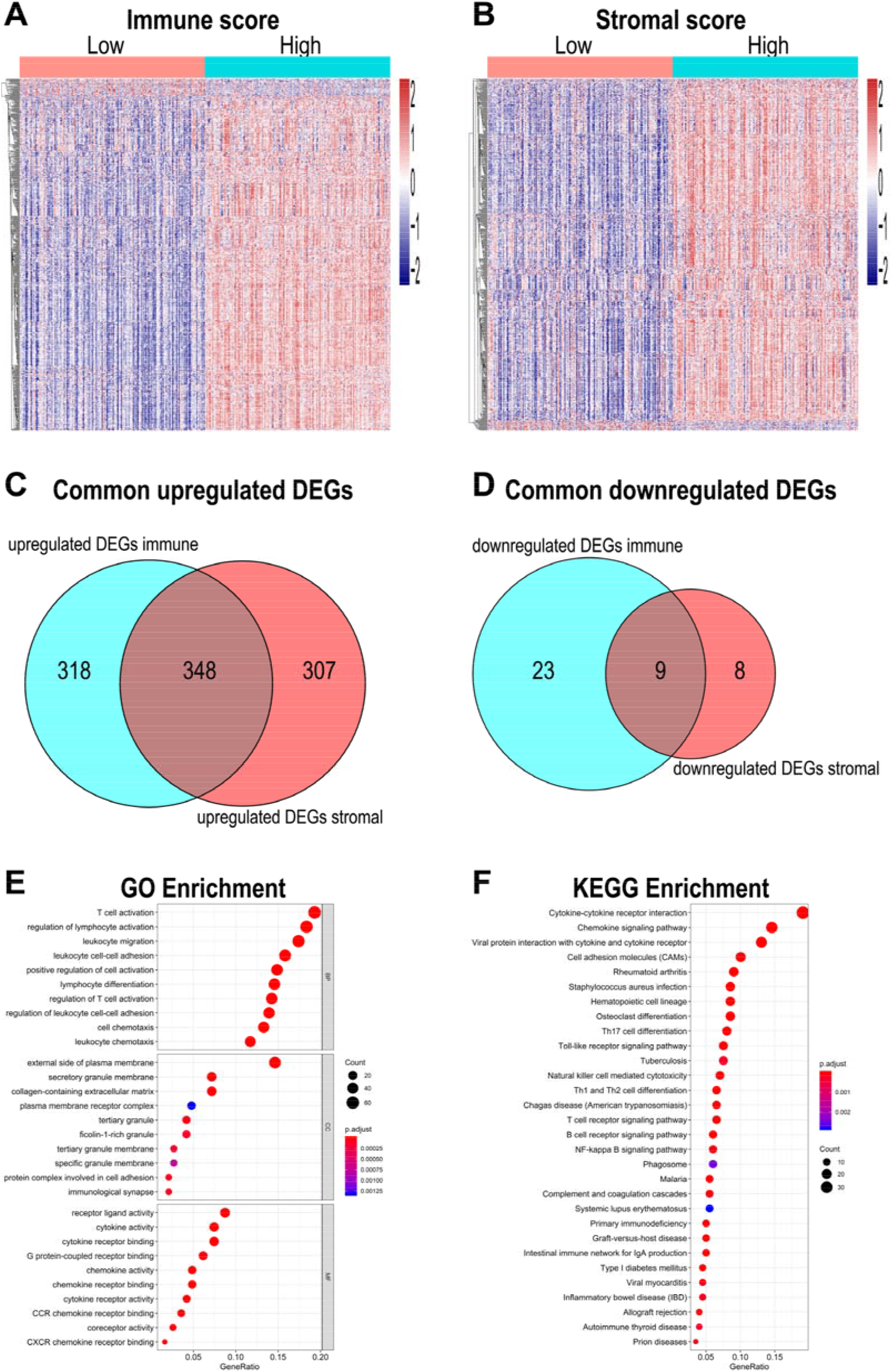
Comparison of gene expression profile with immune scores and stromal scores in lung adenocarcinoma. Heatmaps were drawn based on the average method and correlation distance measurement method. Genes with higher expression are shown in red, lower expression are shown in blue, genes with same expression level are in white. (**A**) Heatmap of the DEGs of immune scores of top half (high score) vs. bottom half (low score). (Cutoff: |log_2_(fold-change) |> 0.75, FDR < 0.05). (**B**) Heatmap of the DEGs of stromal scores of top half (high score) vs. bottom half (low score). (Cutoff: |log2(fold-change) |> 0.75, FDR < 0.05). (**C, D**) Venn diagrams showing the number of common upregulated (**C**) or downregulated (**D**) DEGs in stromal and immune score groups. (**E, F**) Top ten GO terms (**E**) and top thirty KEGG pathway (**F**) enrichments of common DEGs (FDR < 0.05). DEG: differential gene expression; GO: Gene Ontology; BP: biological process; CC: cellular component; MF: molecular function; KEGG: Kyoto Encyclopedia of Genes and Genomes; FDR: false discovery rate.

To illustrate the potential function existed among the 357 common DEGs, we conducted GO enrichment analysis. Top GO terms identified included the T cell activation, the regulation of lymphocyte activation, the external side of plasma membrane, the secretory granule membrane, the receptor ligand activity, and the cytokine activity (**Figure 3E**). Then, we did KEGG pathway enrichment analysis. As plotted in **Figure 3F**, enrichment of common DEGs was major observed for the Cytokine−cytokine receptor interaction, the Chemokine signaling pathway, the Viral protein interaction with cytokine and cytokine receptor, and the Cell adhesion molecules (CAMs).

### Identification of prognostic genes

To identify the potential value of the common DEGs in predicting the overall survival of LUAD patients, we established Kaplan-Meier survival curves. According to the log-rank test (p-value < 0.05), in the common 357 DEGs, 108 genes owned predictive power over overall survival significantly with 103 positively and 5 negatively related (**Supplementary Table S1**). Representative genes are shown in **Figure 4**.

**Figure 4.**
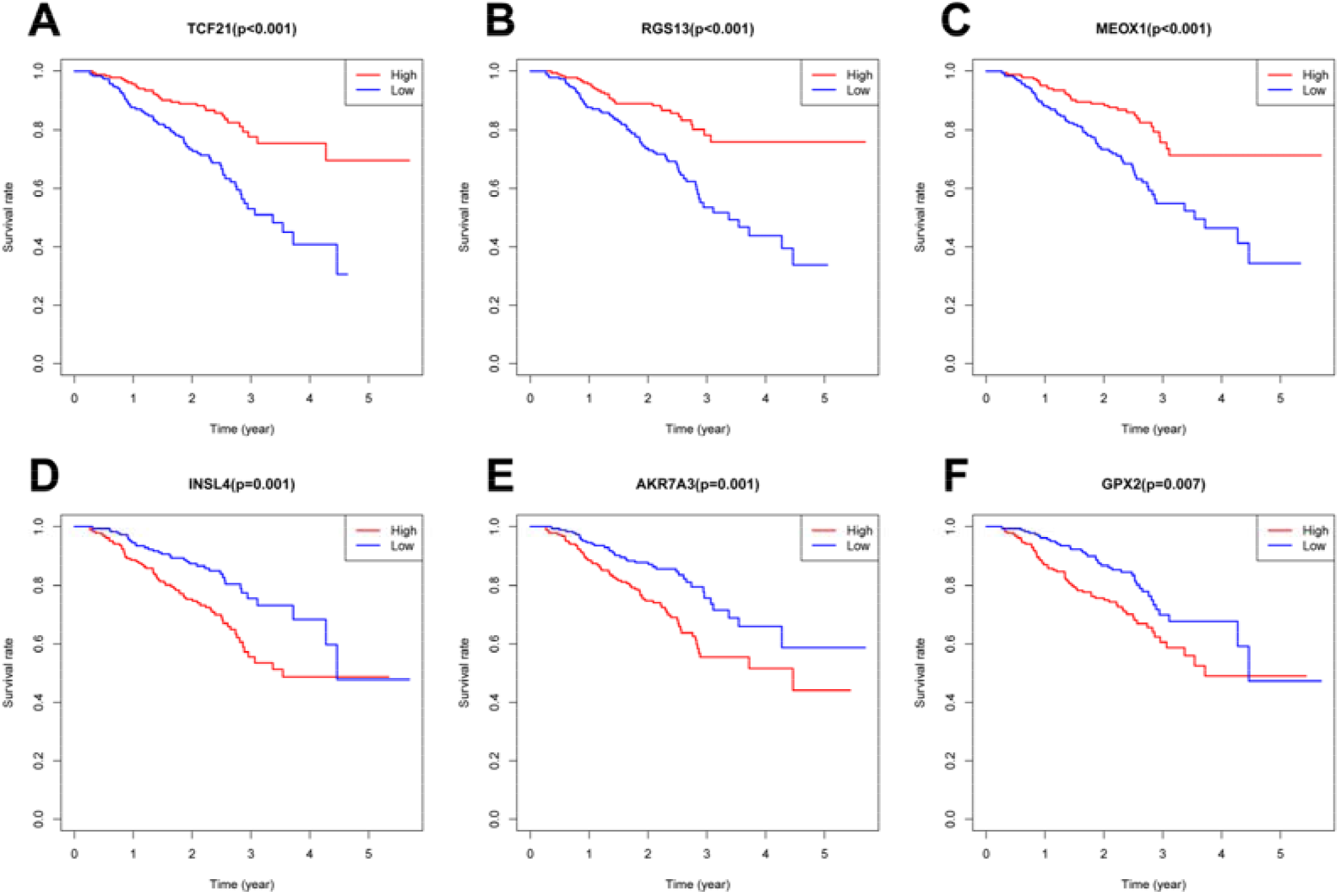
Kaplan–Meier overall survival curves for representative prognostic genes. Comparison of overall survival in the high (**red**) and low (**blue**) gene expression groups. P-value < 0.05 was used to assess differences in the Log-rank test.

### Functional enrichment analysis and protein-protein interactions construction of prognostic genes

GO and KEGG enrichment was conducted to deeper assess the biological function of 108 prognostic genes. These genes involved largely in several biological process (BP), including the T cell activation and the lymphocyte differentiation. The cellular component (CC) suggested that the prognostic genes were generally involved in the external side of plasma membrane and the secretory granule membrane, while the molecular function (MF) process were mainly related to the DNA−binding transcription activator activity, RNA polymerase II−specific and the carbohydrate binding (**Figure 5A**). The KEGG pathways were mainly enriched for the Cytokine−cytokine receptor interaction, the Hematopoietic cell lineage, the Cell adhesion molecules (CAMs), and the Chemokine signaling pathway (**Figure 5B**).

**Figure 5.**
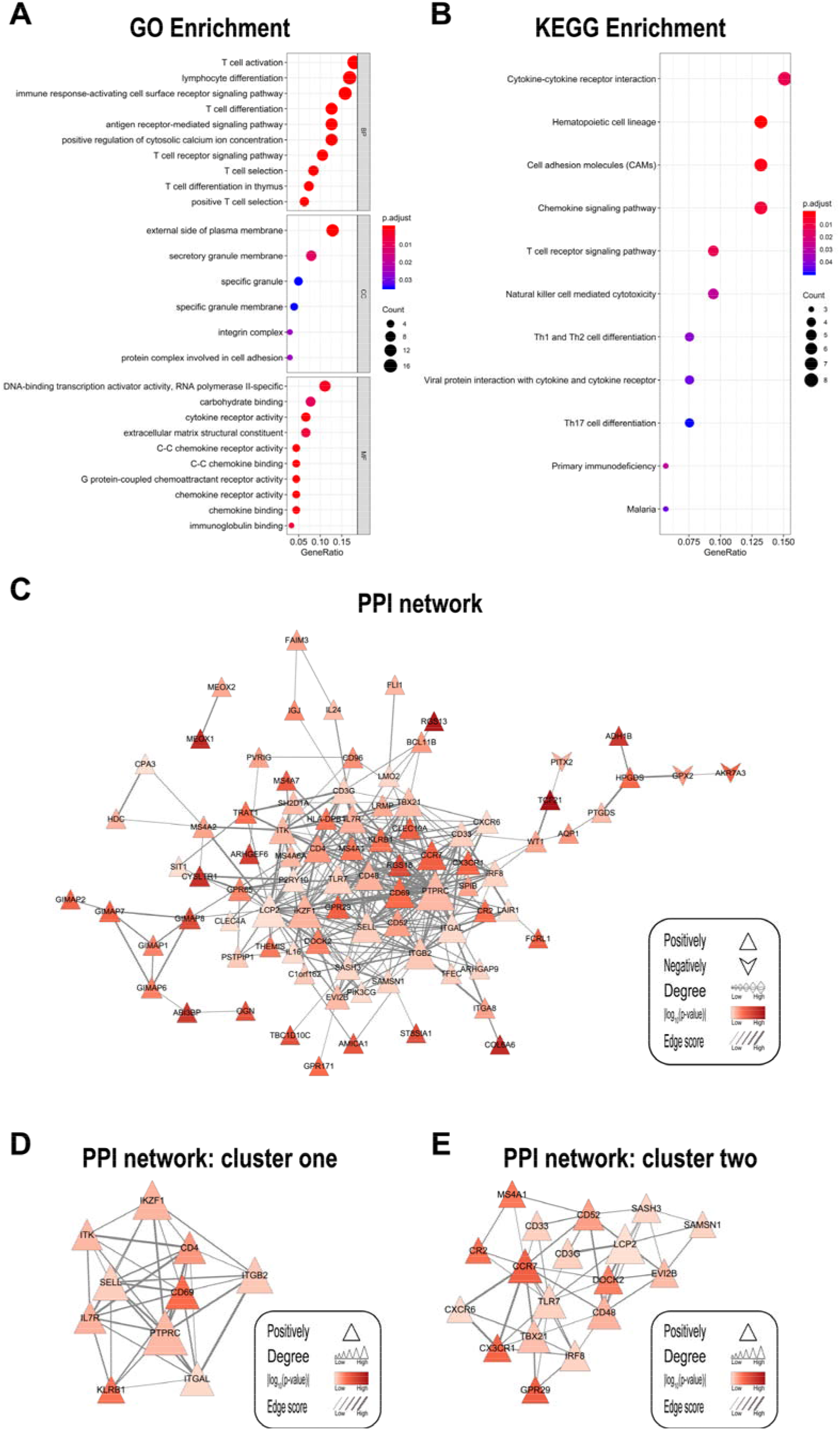
GO and KEGG pathway enrichment and PPI network (with two clusters) of prognostic genes. (**A**) Top ten (If applicable) GO terms with FDR<0.05. (**B**) Top eleven KEGG pathways with FDR<0.05. (**C, D, E**) PPI network (**C**) with two clusters (**D, E**). The “p-value” of the prognostic genes showed in PPIs (**C, D, E**) were gained from the overall survival analysis in GSE72094. “Positively” or “Negatively” meant the gene positively or negatively correlated the overall survival. “Degree” was obtained by the “cytoHubba” plugin from Cytoscape. “Edge score” (combine score) meant the strength between two-gene and was obtained from STRING online tool (https://string-db.org/). GO: Gene Ontology; BP: biological process; CC: cellular component; MF: molecular function; KEGG: Kyoto Encyclopedia of Genes and Genomes; FDR: false discovery rate; PPI: protein-protein interaction.

In order to better understand the interaction among the prognostic genes, we constructed a PPI network via Cytoscape software and STRING online tool. After hiding disconnected nodes, there were 88 nodes and 333 edges in this network (**Figure 5C**). In this network, 85 genes positively and 3 genes negatively correlated with the overall survival of LUAD patients. PTPRC, LCP2, IKZF1, ITGB2, SELL, CD69, ITGAL, CCR7, TLR7, and CD48 were the top ten genes in this network sorted by degree. TCF21, RGS13, MEOX1, COL6A6, CYSLTR1, ADH1B, ABI3BP, ARHGEF6, RGS18, and GIMAP8 owned the top ten predictive ability in this network for overall survival. The interaction between ITGB2 and ITGAL was the tightest in this PPI network, with a combine score of 0.999. The loosest relationship happened between IRF8 and TFEC, with a combine score of 0.4.

Further, we deeply investigated this PPI network using the MCODE plugin in Cytoscape. The network was made up of 3 clusters, of which the top two modules owned ≥10 nodes were plotted (**Figure 5D-E**). In cluster one, PTPRC, IKZF1, ITGB2, SELL, and CD69 hold the top five places sorted by degree, CD69, KLRB1, CD4, IL7R, and IKZF1 owned the top five prognostic ability. While in cluster two, LCP2, CCR7, TLR7, CD48, and TBX21 were the top five degree distribution genes, GPR29, CCR7, CX3CR1, CR2, and MS4A1 were the top five prognostic ability genes.

### Validation in two independent data source

We then validated the 108 prognostic genes using the GSE68465 dataset from the GEO database. P-value< 0.05 in the log-rank test was set as the cutoff. A total of 20 genes were screened, of which 19 genes positively and 1 gene negatively associated with overall survival. Representative Kaplan-Meier survival curves are shown in **Figure 6A-D**. At the same time, we screened these 108 genes in the TIMER, finding 72 genes (70 genes positively, 2 genes negatively) associated with overall survival (cutoff: p-value< 0.05 in the log-rank test). Representative Kaplan-Meier survival curves extracted from the TIMER are shown in **Figure 6E**. After validated in two independent data source, there were 12 genes identified as crucial prognostic genes, all of them were positively associated with overall survival (**Figure 6F**).

**Figure 6.**
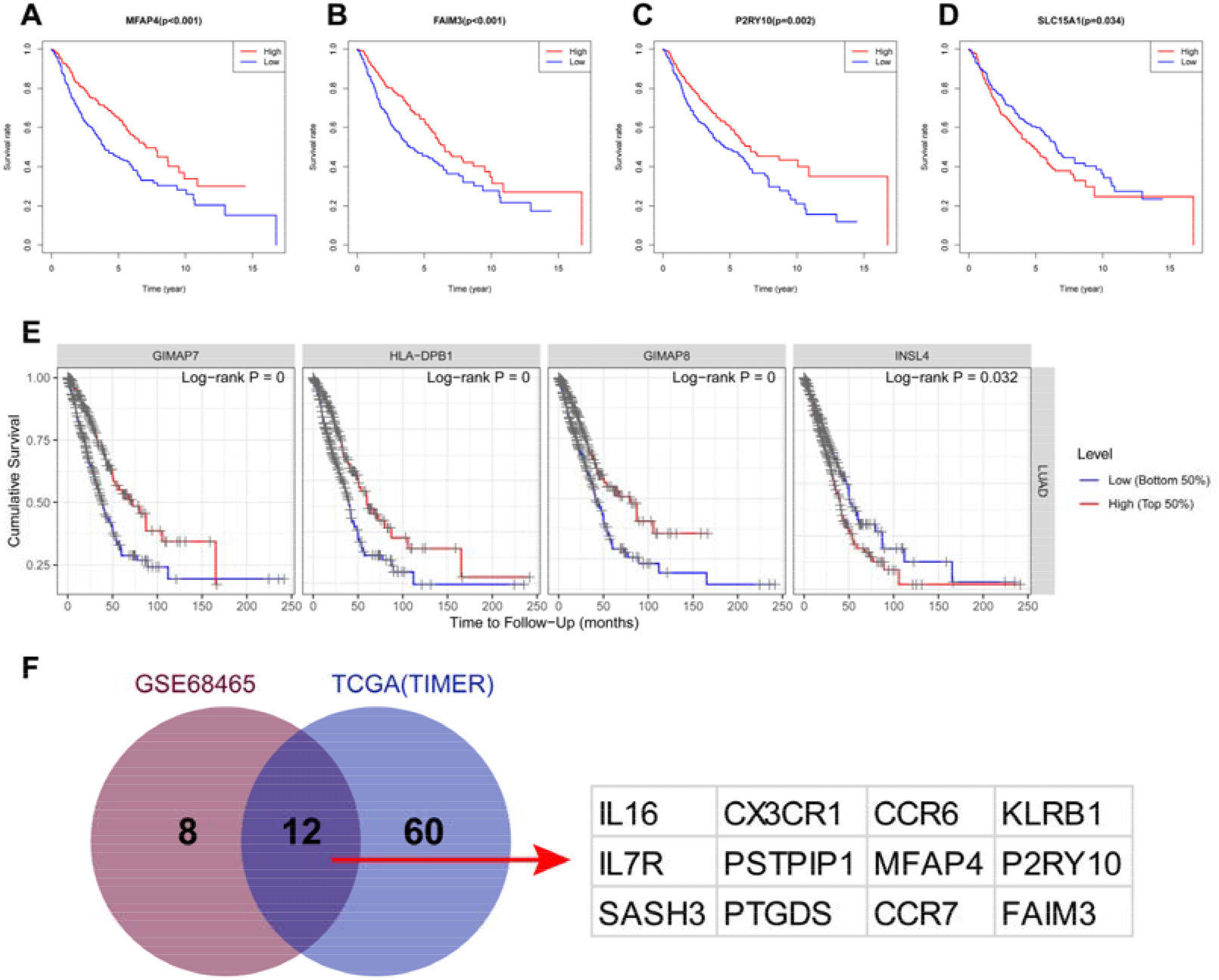
The process of verification in two independent data sources. (**A-D**) Kaplan-Meier overall survival curves of representative genes valid in GSE68465. (**E**) Kaplan-Meier overall survival curves of representative genes valid in TCGA (TIMER). (**F**) Venn diagram based on validation results of two datasets. The exact intersection genes are shown in the right part. P-value < 0.05 was used to assess differences in the Log-rank test.

### Validation of the immune correlates of the crucial prognostic genes

12 crucial prognostic genes are potential immunotherapeutic targets, and their relationships and interactions with immune cells are of great value for further immune-related research. TIMER was applied to verify the correlation between the crucial prognostic genes and immune cell infiltration levels. The correlation analysis results between these 12 genes and CD4 + T cells, CD8 + T cells, B cells, neutrophils, macrophages, and dendritic cells are shown in **Figure 7A**. Graphs of representative genes were plotted in **Figure 7B**. The results showed that although the correlation between the 12 genes and immune cells varies from strong to weak, all correlations were statistically significant.

**Figure 7.**
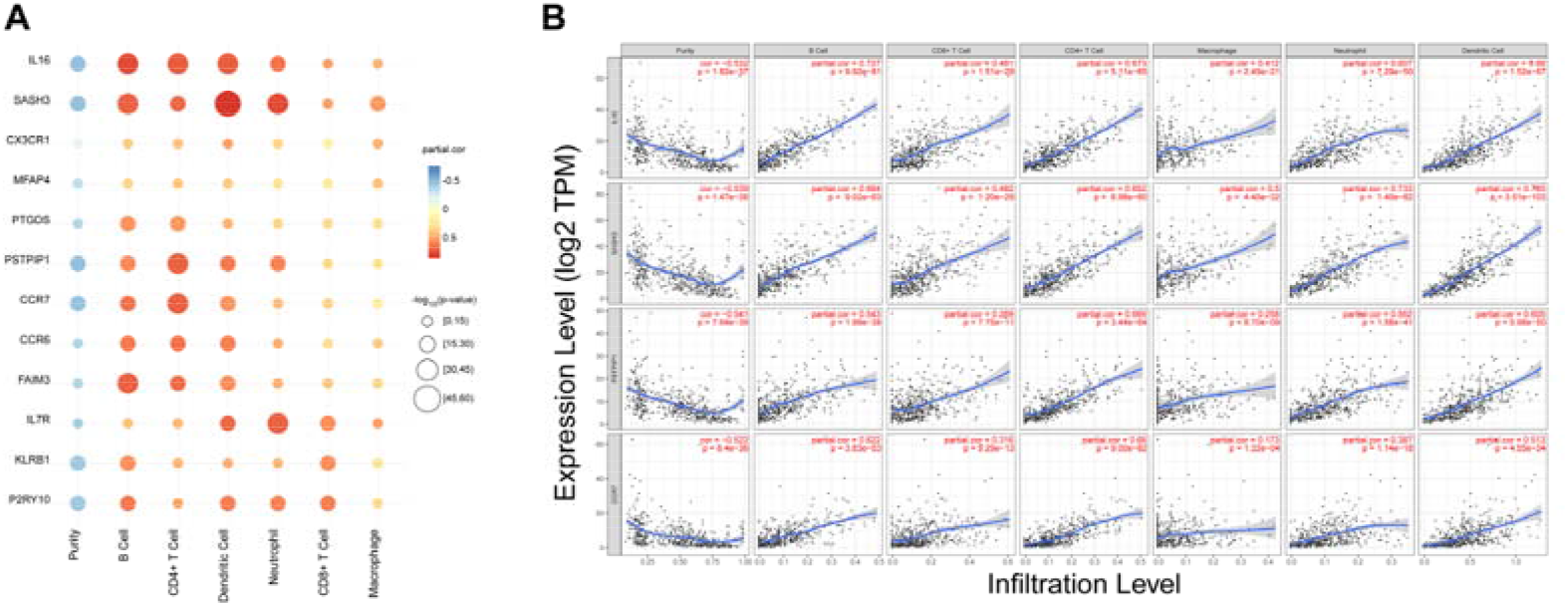
The correlation between the crucial prognostic genes and 6 types of immune cells in lung adenocarcinoma. (**A**) Red represents positive correlation genes and blue represents a negative correlation. The point size represents −log_100_(p-value) and shade of color represents partial.cor (partial correlation coefficient). The x-axis indicates immune cell types, and the y-axis indicates crucial prognostic genes. (**B**) Correlation of representative crucial prognostic genes’ expression with immune infiltration level. Each dot represents a sample, and the blue line represents the relationship between the expression level of each gene and immune cell contents. P-value < 0.05 was considered statistically significant.

## Discussion

In the present study, we attempted to identify tumor microenvironment immune-related genes that contribute to LUAD overall survival. In particular, by comparing global gene expression in cases with high vs. low scores, we obtained 357 common DEGs. Then 108 genes were identified as prognostic genes via overall survival analysis. Importantly, we got 12 genes having crucial prognostic value by validating in two independent data sources. Finally, we employed TIMER to verify the immune correlates of the crucial prognostic genes and found that they all won (**Figure 8**). In addition, the datasets GSE72094 and GSE68465 used in this study have never been applied for tumor microenvironment data mining research before, coupled with the assistance validation of TIMER (based on a large sample of TCGA database) to make our research more novel and rigorous. Our research not only identified the cellular and gene targets of LUAD immunotherapy but also proposed new research ideas for other tumor immunotherapy.

**Figure 8.**
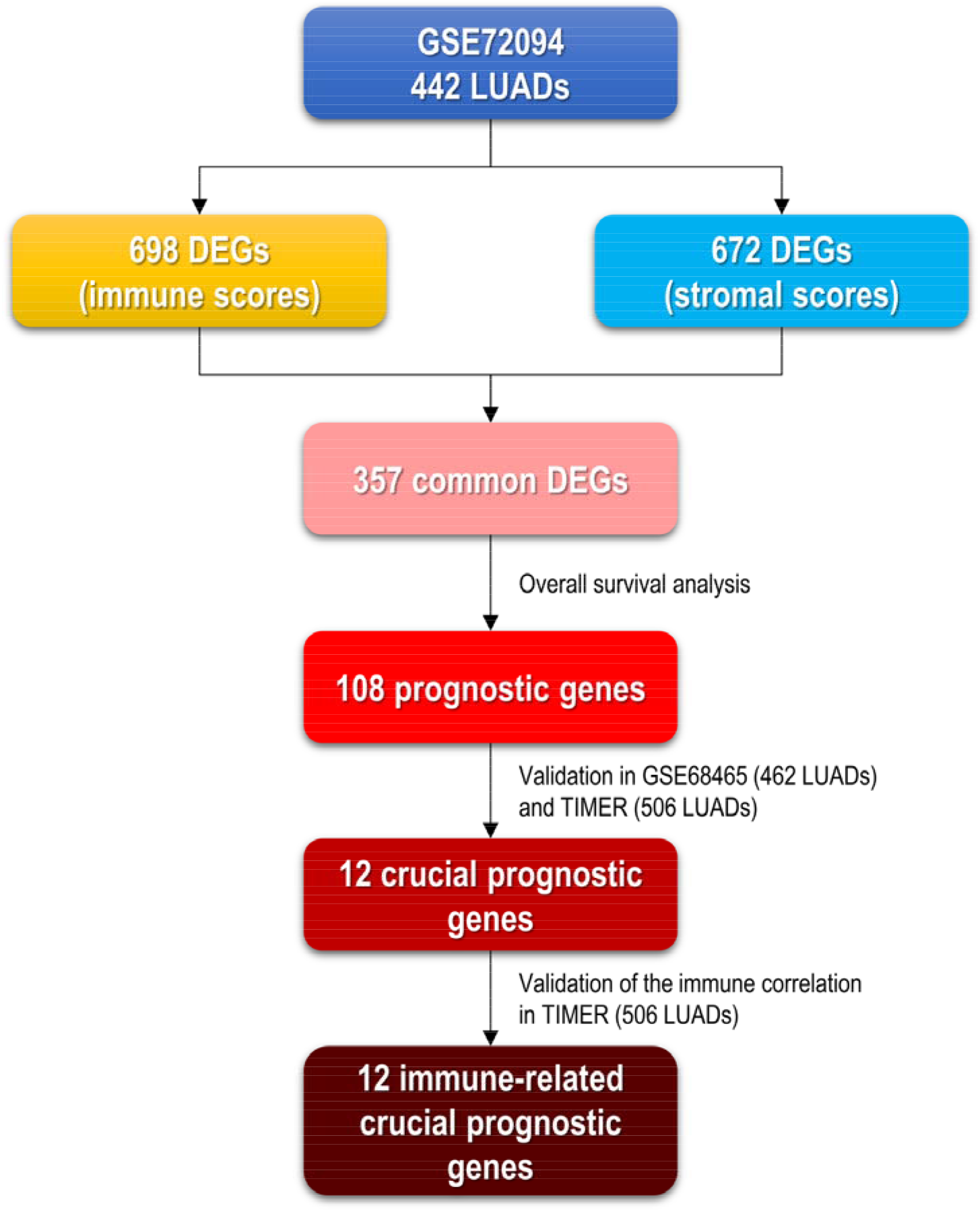
The brief workflow of the present work. DEG: differentially expressed gene; GSE72094, GSE68465: datasets come from Gene Expression Omnibus. LUADs: lung adenocarcinoma cases. TIMER: Tumor IMmune Estimation Resource, https://cistrome.shinyapps.io/timer/.

Firstly, we calculated the immune/stromal scores of LUAD patients based on the ESTIMATE algorithm finding the distributions of immune scores did not vary with gender, age, smoking history, tumor stage, and EGFR status, but vary with the status of KRAS, STK11, and TP53. In the distributions of the stromal scores, only the status of STK11 and TP53 mattered. EGFR is found on the surface of some normal cells and is involved in cell growth. Blocking EGFR may keep cancer cells from growing^26^. KRAS mutations are clear drivers of tumor growth and are characterized by complex biology involving the interaction between mutant KRAS, various growth factor pathways, and tumor suppressor genes^27^. Research shows a gene called STK11, mutated or deleted in a third of non-small cell lung cancer patients, fosters an immunologically “cold” tumor microenvironment, with minimal penetration of tumors by T cells, rendering anti-PD1/PDL1 drugs ineffective^28^. Abnormality of the TP53 gene is one of the most significant events in lung cancers and plays an important role in the tumorigenesis of lung epithelial cells^29^. The relationship between the mutations of these genes and immune/stromal scores can give researchers many hints for further studies.

We also found that reduced immune score predicted a poor prognosis of LUAD. Previous studies have demonstrated that immune cells and stromal cells are important elements active in the tumor microenvironment, affecting LUAD cell survival, proliferation, and therapeutic resistance^30–32^. The crosstalk between tumor and microenvironment influences the inflammatory response: cancer cells interact with both the innate and the adaptive immune system and use immune cells for tumor survival and protection from immunological attacks^33–36^.

In addition, we identified common DEGs from the comparison between low and high immune score / stromal score groups. GO analysis shown these DEGs to be largely enriched in the T cell activation (BP), the external side of plasma membrane (CC), and the receptor ligand activity (MF). Besides, the KEGG pathway enrichment displayed that the DEGs mostly clustered in the Cytokine−cytokine receptor interaction, the Chemokine signaling pathway, and the Viral protein interaction with cytokine and cytokine receptor. Consistent with these results, previous studies have shown that the biological processes of the immune system are critical to the formation of a complicated LUAD tumor microenvironment^10,30,37–39^. In the past few years, the understanding of the immunological characteristics of LUAD has increased, and the development of effective LUAD immunotherapy strategies has drawn wide attention^40–44^.

Overall survival analysis of the common DEGs revealed that 108 genes (30.3%) owned predictive ability over the overall survival of LUAD patients significantly. Moreover, the PPI network and two clusters were constructed to reveal the relationship and function of prognostic genes. Nodes with a high connectivity degree in the PPI network, including PTPRC, LCP2, IKZF1, ITGB2, SELL, CD69, ITGAL, CCR7, TLR7, and CD48 were potentially related to the prognosis of LUAD (**Figure 5C**). Among them, CD69 and IKZF1 were in the top five in both connectivity degree and predictive ability of cluster one (**Figure 5D**), and CCR7 also had the same performance in cluster two (**Figure 5E**). The three genes had indeed caught our attention. CD69 is an early activation marker that is expressed in hematopoietic stem cells, T cells, and many other cell types in the immune system^45^. In addition to its intrinsic value as an activation marker, CD69 is also an important regulator of immune responses^46^. Study had shown that CD69 plays a vital role in antitumor immunity by regulating the exhaustion of tumor-infiltrating T cells^47^. IKZF1 is a key regulator of a complex molecular signature governing immune infiltrate recruitment, making it both a predictor of response to therapy and an excellent candidate for future drug targeting^48^. IKZF1 could enhance the sensitivity of immune Infiltrate recruitment and immunotherapy of solid tumors^48^. In the tumor microenvironment, chemokines and chemokine receptors have essential roles in tumor proliferation, metastasis, and invasion. CCR7 was confirmed to participate in cancer cell metastasis, invasion, and tumor development^49–53^.

In this study, GSE68465 (462 LUAD cases) and TIMER (506 LUAD cases) were used for validation. 12 genes were identified as crucial prognostic genes of LUADs. Tumor-infiltrating immune cells play an important role in promoting or inhibiting tumor growth. As an integral part of the tumor microenvironment, tumor-infiltrating immune cells exist in all stages of cancer and play a vital role in shaping the development of tumors^54^. To study the effect of immune cell infiltration on the tumor microenvironment on the prognosis of LUAD patients, we used the TIMER to calculate the extent of infiltrations of CD4 + T cells, CD8 + T cells, B cells, neutrophils, macrophages, and dendritic cells and correlated the data with the expression of the identified crucial prognostic genes. Although the results showed that the associations between the 12 genes and immune cells vary from strong or weak, they are all statistically significant, indicating that all 12 genes are associated with the infiltrations of these 6 main types of immune cells (**Figure 7**). In the graph plotted in **Figure 7A**, IL16, SASH3, PSTPIP1, CCR7, CCR6, FAIM3, and IL7R seemed to attract more kinds of immune cells infiltrated than the rest. From another point of view, B cells, CD4+ T cells, dendritic cells, and neutrophils involved infiltrating in more genes than other types of immune cells.

## Conclusion

In summary, from functional enrichment analysis of a GEO dataset applied by ESTIMATE algorithm-based stromal/immune scores, we obtained a list of genes related to the tumor microenvironment. After the overall survival analysis and two independent validation, we got 12 crucial prognostic genes having the ability to illustrate the prognosis of LUAD patients. Amazingly, all identified crucial prognostic genes shown significantly correlated with the infiltration of CD4 + T cells, CD8 + T cells, B cells, neutrophils, macrophages, and dendritic cells. It would be interesting to test whether this set of genes or targeting these 6 types of immune cells, when combined, can provide more influence on the outcome of LUAD than an individual gene or targeting. Further research on these immune-related genes may reveal a potential association between tumor microenvironment and LUAD prognosis in a novel and comprehensive way, providing a more theoretical basis for the development of new LUAD prognostic biomarkers and immunotherapy.

## Data Availability Statement

Publicly available datasets were analyzed in this study. These data can be found here: TCGA: https://portal.gdc.cancer.gov/; GEO: https://www.ncbi.nlm.nih.gov/geo/.

## Ethics Statement

Ethical review and approval were not required for the study on human participants in accordance with the local legislation and institutional requirements. Written informed consent for participation was not required for this study in accordance with the national legislation and the institutional requirements.

## Acknowledgments

Chao Ma and Huan Luo thank Zhengzhou University Overseas Virtual Research Institute for financing support studying abroad. Chao Ma thanks to China Scholarship Council (No. 201708410121) for its financial supporting. We acknowledge support from the German Research Foundation (DFG) and Charité – Universitätsmedizin Berlin.

## Author Contributions

Chao Ma and Huan Luo organized and wrote the manuscript. Chao Ma designed and produced the figures. Huan Luo contributed to the literature search for the manuscript. Jing Cao revised the manuscript. All authors reviewed the manuscript and approved the manuscript for publication.

